# Comparative genomics of *Nocardia seriolae* reveals recent importation and subsequent widespread dissemination in mariculture farms in South Central Coast, Vietnam

**DOI:** 10.1101/2021.11.30.470482

**Authors:** Cuong T. Le, Erin P. Price, Derek S. Sarovich, Thu T.A Nguyen, Daniel Powell, Hung Vu-Khac, Ipek Kurtböke, Wayne Knibb, Shih-Chu Chen, Mohammad Katouli

**Author notes:** **Corresponding author:** A/Prof Mohammad Katouli, Genecology Research Centre and School of Science, Technology and Engineering, University of the Sunshine Coast, Locked Bag 4, Maroochydore BC, Queensland, 4558, Australia. **Repositories**: The Illumina data and genome assemblies for the seven Vietnamese *N. seriolae* genomes generated in this study are available at NCBI BioProject PRJNA551736.

## Abstract

Between 2010 and 2015, nocardiosis outbreaks caused by *Nocardia seriolae* affected many permit farms throughout Vietnam, causing mass fish mortalities. To understand the biology, origin, and epidemiology of these outbreaks, 20 *N. seriolae* strains collected from farms in four provinces in the South-Central Coast of Vietnam, along with two Taiwanese strains, were analysed using genetics and genomics. Pulsed-field gel electrophoresis identified a single cluster amongst all Vietnamese strains that was distinct from the Taiwanese strains. Like the PFGE findings, phylogenomic and single-nucleotide polymorphism (SNP) genotyping analyses revealed that all Vietnamese *N. seriolae* strains belonged to a single, unique clade. Strains fell into two subclades that differed by 103 SNPs, with almost no diversity within clades (0-2 SNPs). There was no association between geographic origin and subclade placement, suggesting frequent *N. seriolae* transmission between Vietnamese mariculture facilities during the outbreaks. Vietnamese strains shared a common ancestor with strains from Japan and China, with the closest strain, UTF1 from Japan, differing by just 217 SNPs from the Vietnamese ancestral node. Draft Vietnamese genomes range from 7.55-7.96 Mbp in size, have an average G+C content of 68.2%, and encode 7,602-7,958 predicted genes. Several putative virulence factors were identified, including genes associated with host cell adhesion, invasion, intracellular survival, antibiotic and toxic compound resistance, and haemolysin biosynthesis. Our findings provide important new insights into *N. seriolae* epidemiology and pathogenicity and will aid future vaccine development and disease management strategies, with the ultimate goal of nocardiosis-free aquaculture.

## 1. Introduction

The genus *Trachinotus*, of the family Carangidae, comprises a group of marine, medium-sized, migratory, pelagic finfish that are widely distributed in subtropical and tropical waters worldwide (Berry and Iversen, 1967, Finucane, 1969). Many members, such as *T. carolinus*, *T. blochii, T. ovatus,* and *T. falcatus*, are of great economic importance for fisheries and aquaculture sectors in America and Asia due to high quality-meat, fast growth, high market price, and strong adaptability to a variety of captive environments (Muller et al., 2002, McMaster et al., 2003, Tutman et al., 2004, Klinkhardt and Myrseth, 2007, Juniyanto et al., 2008). In Asia, the farming of permit fish, particularly the snub nose permit, *T. falcatus*, has commercially taken place in ponds, raceways, and floating sea cages in both brackish and sea waters. Since 2010, Asian mariculture farms have produced over 2 million tonnes of fish meat, significantly contributing to food security, poverty alleviation, and economic growth of the region (FAO, 2021). However, the shortage of quality seed stock and the risk of fish disease outbreaks in several countries are key obstacles and challenges for the sector’s sustainable development.

*T. falcatus* fingerlings were first imported into Vietnam from Taiwan and China in the 2000s and have quickly gained popularity, with permit fish now the third largest group of commercially cultured marine fish after seabass and grouper. However, high mortality rates of *T. falcatus* weighing between 5 and 350 g (6 -45 cm in length) emerged in 2010 during an epizootic event that affected sea cage farms in Khánh Hòa province, in the South-Central Coast region of Vietnam. Since this initial outbreak, large-scale outbreaks have occurred at several other farming sites in southern and central parts of the country (Nguyen et al., 2012, Vu-Khac et al., 2016). Infected fish showed clinical signs of nocardiosis such as lethargy, skin blisters, ulcers, and multiple yellowish to whitish nodules affecting both internal and external organs. Based on analyses of 16S rDNA sequences and biochemical characteristics, the bacterial pathogen, *Nocardia seriolae*, was confirmed as the causative agent (Vu-Khac et al., 2016); however, the origin of *N. seriolae* affecting Vietnamese permit fish farms has not yet been identified.

*N. seriolae* is a Gram-positive, branching, filamentous intracellular bacterium of the family Nocardiaceae that was initially described as *N. kampachi* in farmed yellowtail, *Seriola quinqueradiata*, by Kariya et al. (1968) following large outbreaks in Mie Prefecture, Japan. An estimated loss of approximately 260 tonnes of cultured yellowtails due to the disease was recorded in 1989 (Kusuda and Salati, 1993). Nocardiosis has also impacted several other important fish species within the Japanese aquaculture industry such as amberjack (*Seriola dumerili*), Japanese flounder (*Paralichthys olivaceus*), and chub mackerel (*Scomber japonicas*)*. N. seriolae* has subsequently been documented in Taiwan, China, Korea, USA, and Mexico, where high mortalities and associated economic losses due to nocardiosis having been reported in freshwater and marine fish species in both cultured and wild populations (Kudo et al., 1988, Chen et al., 1989, Chen and Tung, 1991, Chen et al., 2000, Huang, 2004, Park et al., 2005, Shimahara et al., 2008, Shimahara et al., 2009, Cornwell et al., 2011, Kim et al., 2018, Del Rio-Rodriguez RE, 2021). Despite causing significant economic losses in fish aquaculture worldwide, there are currently no effective measures against nocardiosis.

Four complete and nine draft *N. seriolae* genome sequences are publicly available as of 16Aug21, representing isolates retrieved from Japan, South Korea, and China (Imajoh et al., 2015, Xia et al., 2015, Imajoh et al., 2016, Yasuike et al., 2017, Han et al., 2018). These genomes have provided important insights into *N. seriolae* epidemiology, transmission, pathogenesis, and infection control strategies; however, isolates from other nocardiosis-prevalent regions such as Vietnam have not yet been examined, leaving major gaps in our understanding of this devastating infectious disease. In the current study, we sequenced the entire genomes of seven *N. seriolae* isolates isolated from different permit fish farm locations across Vietnam and compared them with the 12 previously genome-sequenced *N. seriolae* isolates, allowing a comparison of isolates spanning a decade time scale and from a variety of sources and geographic locations. Using this information, we developed two novel single-nucleotide polymorphism (SNP)-based PCR assays to rapidly differentiate Viet and non-Viet strains, and strains representing the two Vietnamese clades. We also characterised potential virulence factors and antimicrobial/toxin resistance determinants to gain insights into pathogenicity and survival mechanisms. Finally, we functionally annotated our *N. seriolae* genomes to determine whether differences in gene content might contribute to physiological variability among isolates.

## Methods

### Bacterial strains

Due to a ban on *N. seriolae* culture importation into Australia, all live culture work was carried out in laboratories at Institute of Aquaculture, Nha Trang University, Vietnam (for Vietnamese strains) and the Department of Veterinary Medicine, College of Veterinary Medicine, National Pingtung University of Science and Technology, Pingtung, Taiwan.

Twenty-two *N. seriolae* strains isolated from fish were examined in this study, comprising 20 from Vietnam and two from Taiwan. Vietnamese strains were isolated from cultured permit fish (*T. falcatus*) (31.0 – 85.8g) during nocardiosis outbreaks occurring between 2014 and 2015 in four provinces (Phú Yên, Khánh Hòa, Ninh Thuận, and Vũng Tàu) in the South Central Coast region, and the Taiwanese strains were isolated from largemouth bass (*Micropterus salmoides)* and mullet (*Mugil cephalus)* in 2007 (Fig. 1 and Table 1). Isolates were confirmed as *N. seriolae* based on morphological observations, Ziehl-Neelsen staining (Fig. 2), 16S sequencing, and biochemical characteristics (Vu-Khac et al., 2016). The 20 Vietnamese strains were subject to pulsed-field gel electrophoresis (PFGE) analyses, of which seven isolates were selected for whole-genome sequencing (WGS) to enable more detailed genetic analyses. All 22 isolates were tested using our SNP genotyping assays.

**Fig. 1.**
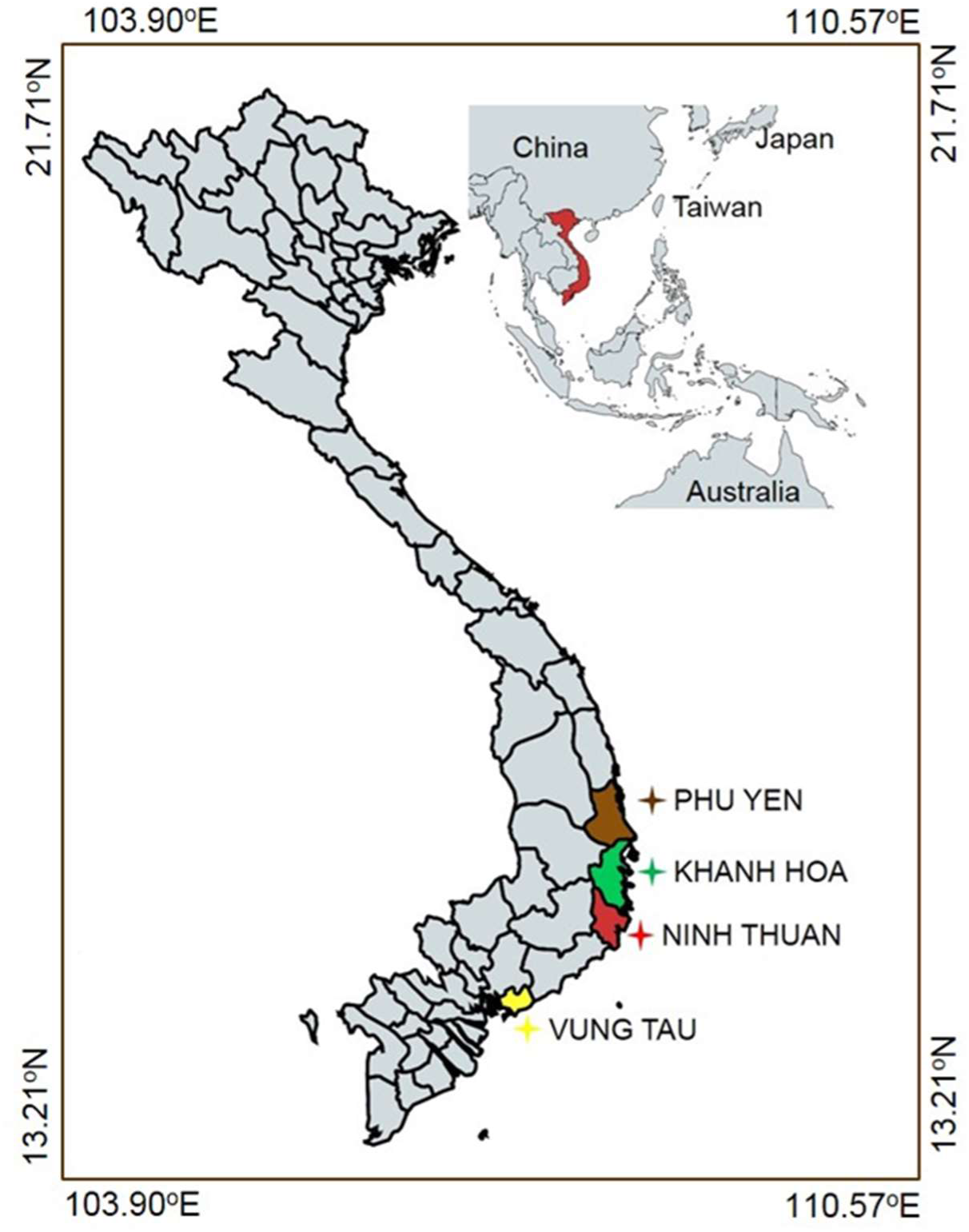
Four Vietnamese provinces where *Nocardia seriolae* isolates were collected from infected permit fish (*Trachinotus falcatus*).

**Fig. 2.**
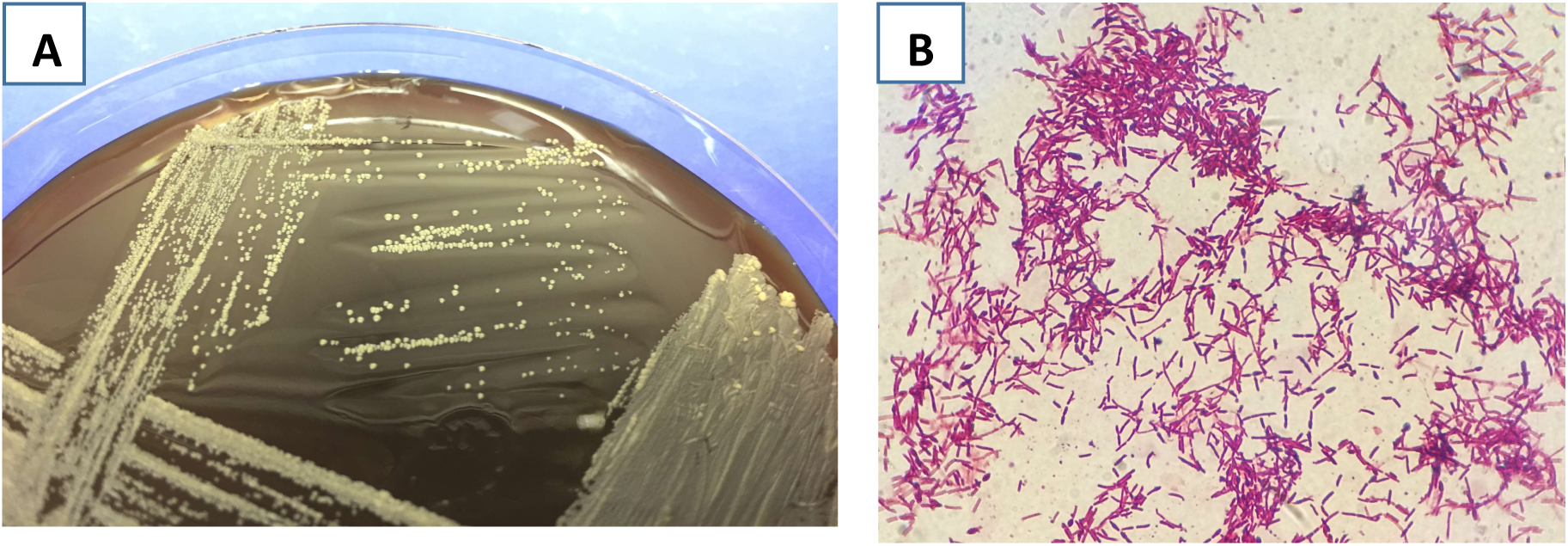
Morphology of *Nocardia seriolae* isolated from Vietnam mariculture farms. (A) Chalky white non-hemolytic colonies of *N. seriolae* on sheep blood agar (3 week-old culture); and (B) Ziehl–Neelsen stained *N. seriolae*, showing purple red, filamentous branching bacteria.

**Table 1.** Nocardia seriolae strains collected in this study, their *Ase*I and *Xba*I pulsed-field gel electrophoresis profiles, and their single-nucleotide polymorphism (SNP) genotypes. *S1, non-Vietnamese SNP genotype; S2, Vietnamese SNP genotype; C1, Vietnam Clade 1; C2, Vietnam Clade 2

| Country | Strain | Fish species | Host tissue | Origin | Collection date | *AseI* | *XbaI* | SNP genotype* |
| --- | --- | --- | --- | --- | --- | --- | --- | --- |
| Taiwan | 96127 | *Micropterus salmoides* |  | Taiwan | 2007 | A1 | X1 | S1 |
| Taiwan | 96994 | *Mugil cephalus* |  | Taiwan | 2007 | A4 | X5 | S1 |
| Vietnam | KH_11 | *Trachinotus falcatus* | Muscle | Khánh Hòa, Vietnam | Mar 2014 | NsA2 | NsX3 | S2C1 |
| Vietnam | KH_14 | *Trachinotus falcatus* | Spleen | Khánh Hòa, Vietnam | Apr 2014 | NsA1 | NsX1 | S2C2 |
| Vietnam | KH_15 | *Trachinotus falcatus* | Kidney | Khánh Hòa, Vietnam | May 2014 | NsA1 | NsX5 | S2C1 |
| Vietnam | KH_17 | *Trachinotus falcatus* | Spleen | Khánh Hòa, Vietnam | Mar 2014 | NsA1 | NsX3 | S2C1 |
| Vietnam | KH_21 | *Trachinotus falcatus* | Kidney | Khánh Hòa, Vietnam | Apr 2014 | NsA2 | NsX3 | S2C2 |
| Vietnam | NT_01 | *Trachinotus falcatus* | Muscle | Ninh Thuận, Vietnam | Apr 2014 | NsA3 | NsX5 | S2C2 |
| Vietnam | NT_02 | *Trachinotus falcatus* | Spleen | Ninh Thuận, Vietnam | Apr 2014 | NsA3 | NsX2 | S2C1 |
| Vietnam | NT_03 | *Trachinotus falcatus* | Liver | Ninh Thuận, Vietnam | Apr 2014 | NsA5 | NsX1 | S2C2 |
| Vietnam | NT_50 | *Trachinotus falcatus* | Spleen | Ninh Thuận, Vietnam | Apr 2014 | NsA2 | NsX3 | S2C2 |
| Vietnam | PY_22 | *Trachinotus falcatus* | Spleen | Phú Yên, Vietnam | Apr 2014 | NsA4 | NsX1 | S2C1 |
| Vietnam | PY_23 | *Trachinotus falcatus* | Muscle | Phú Yên, Vietnam | Apr 2014 | NsA9 | NsX1 | S2C1 |
| Vietnam | PY_30 | *Trachinotus falcatus* | Liver | Phú Yên, Vietnam | Apr 2014 | NsA8 | NsX1 | S2C2 |
| Vietnam | PY_31 | *Trachinotus falcatus* | Bone | Phú Yên, Vietnam | Apr 2014 | NsA10 | NsX4 | S2C1 |
| Vietnam | PY_35 | *Trachinotus falcatus* | Spleen | Phú Yên, Vietnam | Apr 2014 | NsA7 | NsX1 | S2C2 |
| Vietnam | PY_37 | *Trachinotus falcatus* | Spleen | Phú Yên, Vietnam | Apr 2014 | NsA6 | NsX1 | S2C2 |
| Vietnam | PY_39 | *Trachinotus falcatus* | Spleen | Phú Yên, Vietnam | Apr 2014 | NsA7 | NsX1 | S2C2 |
| Vietnam | PY_40 | *Trachinotus falcatus* | Kidney | Phú Yên, Vietnam | Apr 2014 | NsA6 | NsX1 | S2C1 |
| Vietnam | VT_45 | *Trachinotus falcatus* | Spleen | Vũng Tàu, Vietnam | Jun 2015 | NsA10 | NsX3 | S2C1 |
| Vietnam | VT_61 | *Trachinotus falcatus* | Spleen | Vũng Tàu, Vietnam | Jun 2015 | NsA11 | NsX1 | S2C1 |
| Vietnam | VT_62 | *Trachinotus falcatus* | Liver | Vũng Tàu, Vietnam | Jun 2015 | NsA12 | NsX1 | S2C2 |

Isolates were preserved in Brain Heart Infusion (BHI, Difco, Sparks, MD, USA) broth mixed with 25% (v/v) glycerol and stored at -80°C. For culturing, strains were grown in BHI broth at 28°C for five days, with orbital shaking at 150 rpm. For DNA extraction, 0.3 mL of bacterial cells were pelleted at 6000 x *g* at 4°C for 5 min and washed twice with 1X sterile phosphate-buffered saline. To test for haemolytic reaction, *N. seriolae* colonies grown in BHI broth were streaked onto 5% (v/v) sheep blood agar and incubated at 28°C for three weeks (Fig. 2).

### PFGE typing

PFGE was performed using 50 U *Xba*I or *Ase*I (New England BioLabs, Ipswich, MA, USA) as previously described (Shimahara et al., 2009). The type strain, *N. seriolae* BCRC 13745 (JCM 3360; isolated from the spleen of farmed yellowtail in Nagasaki Prefecture, Japan, *ca.* 1974) (Kudo et al., 1988), was included for comparative purposes. Gels of DNA fragments were analysed using GelCompar II software version 6.5 (Applied Maths, Kortrijk, Belgium). Gel bands were automatically assigned by the software and were checked and corrected manually. Only clearly resolved bands were considered for further analysis. A dendrogram was constructed using an unweighted pair group method with arithmetic mean (UPGMA) approach and the Dice similarity coefficient, with band optimisation and band position tolerances of 1.0%. Isolates that showed similarity between the banding profiles of ≥80% (fewer than six bands of difference) were defined as indistinguishable or clonally related, whereas patterns with <80% similarity (six or more bands of difference) represented different clusters of unrelated strains (Tenover et al., 1995b, Calvez et al., 2015b).

### DNA extraction

Total genomic DNA of bacterial isolates was extracted using the Wizard® Genomic DNA Purification Kit (Promega, Madison, WI, USA) as per the manufacturer’s instructions. DNA was checked for sterility and shipped to the University of the Sunshine Coast, Queensland, Australia. Quantity and purity of extracted DNA were assessed using a NanoDrop 2000 (Thermo Scientific, Scoresby, VIC, Australia) and 1% gel electrophoresis. DNA for Illumina whole-genome sequencing was submitted on dry ice to the Australian Genome Research Facility (AGRF; North Melbourne, VIC, Australia).

### WGS and comparative genomic analyses

NextEra DNA Flex Illumina libraries for seven Vietnamese *N. seriolae* isolates were sequenced in four lanes of a single flowcell on the NextSeq 500 platform (Illumina, San Diego, CA, USA), to produce 150 bp paired reads at an average depth of ∼ 390× (range: 326 to 433×). Raw read quality was assessed with FastQC v0.11.5 (http://www.bioinformatics.babraham.ac.uk/projects/fastqc/). These seven genomes are available on the Sequence Read Archive database under BioProject PRJNA551736. Twelve publicly available genome assemblies (strains EM150506, CK-14008, HSY-NS01, HSY-NS02, MH196537, N-2927, NBRC 15557, NK201610020, SY-24, U-1, UTF1, and ZJ0503, corresponding to GenBank assembly references ASM186585v1, ASM188553v1, ASM301359v1, ASM366707v1, ASM1411730v1, ASM58371v2, ASM799071v1, ASM1520982v1, ASM209393v1, ASM119293v1, ASM235603v1, and ASM76316v1, respectively) were converted to simulated Illumina reads using ART v2016.06.05 (Huang et al., 2012) prior to analysis. EM150506, the largest complete *N. seriolae* genome (GenBank accession number CP017839.1) (Han et al., 2018), was used as the reference sequence for read mapping and gene annotation. Biallelic, orthologous SNPs from the 19 *N. seriolae* genomes were identified using the default settings of SPANDx v3.2 (Sarovich and Price, 2014), which integrates Burrows-Wheeler Aligner (Li and Durbin, 2009), Sequence Alignment/Map (SAM) tools (Li et al., 2009), BEDTools (Quinlan et al., 2010), VCFtools (Danecek et al., 2011), Picard Tools (http://broadinstitute.github.io/picard) and Genome Analysis Tool Kit (Mckenna et al., 2010) into a single pipeline.

Using the SPANDx SNP matrix (Data S1), a maximum parsimony phylogenomic tree was constructed by Phylogenetic Analysis Using Parsimony (PAUP*) v4.0a168 software (Swofford, 1998), with trees visualised using FigTree v1.4.0 (http://tree.bio.ed.ac.uk/software/figtree/). Variant annotation was carried out using SnpEff (Cingolani et al., 2012). To determine similarity among *N. seriolae* genomes, and to check for potential rearrangements, contigs in all genome assemblies were oriented and arranged against the reference genome using MAUVE v2.3.1 (Darling et al., 2004). BLAST Ring Image Generator (BRIG) (Alikhan et al., 2011) was subsequently used to visualise genome relatedness and structural variation.

### SNP genotyping

The SPANDx SNP matrix was used to identify SNPs that: i) distinguished Vietnamese from non-Vietnamese *N. seriolae* strains (217 SNPs; SNP1 assay), and: ii) differentiated the two Vietnamese clades (103 SNPs; SNP2 assay). We selected SNPs at positions 60409 and 587171 in EM150506 for SNP1 and SNP2 assay design, respectively (Data S1). SYBR green-based mismatch amplification mutation assay (SYBR-MAMA) real-time PCRs were developed to permit rapid genotyping of all strains from this study against these two SNPs. SYBR-MAMA, also known as allele-specific PCR or amplification-refractory mutation system, exploits the differential 3’ amplification efficiency of *Taq* polymerase in real-time via allele-specific primers targeting each SNP allele at their ultimate 3’-end (Germer et al., 2000). SYBR­MAMA has been used for SNP genotyping in many bacteria (Birdsell et al., 2012, Price et al., 2010) due to its low cost and simplicity. Each SNP assay consisted of one common primer and two allele-specific primers, matching either the non-Viet allele or the Viet allele for the SNP1 assay, and the Viet Clade 1 allele or Viet Clade 2 allele for the SNP2 assay (Table 2). The same destabilizing mismatch (A for SNP1 and G for SNP2) was incorporated at the penultimate (-2) 3’ base of both allele-specific primers to increase allele specificity (Hézard et al., 1997). Cycles-to-threshold (CT) values for each allele-specific reaction were used to determine the SNP genotype for each strain via a change in CT value (ΔCT).

**Table 2.**
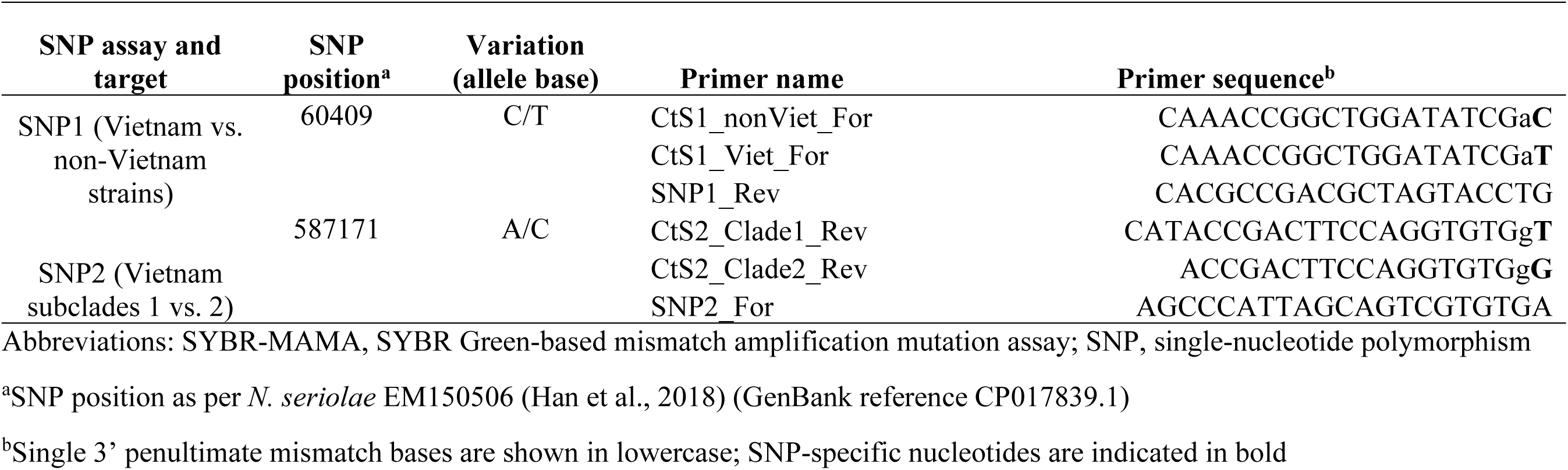
Primer sequences of SYBR-MAMA assays designed in this study for the differentiation of Vietnamese *Nocardia seriolae* strains

To validate SNP genotypes for our newly developed assays, we first established the reference ΔCT values for each assay by running against the two Taiwanese and seven genome-sequenced Vietnamese strains. Assays were then tested against the 13 remaining Vietnamese isolates to determine their genotypes. For each PCR run, control DNA samples representing the matching and non-matching allele genotypes were used as positive controls, and at least two no-template controls were included. SYBR-MAMAs contained 1 μL of target DNA template at ∼1ng/μL, 0.2 µM allele-specific primer, 0.2 µM common primer (Macrogen, Inc., Geumcheon-gu, Seoul, Republic of Korea), 1X Platinum™ SYBR™ Green qPCR SuperMix-UDG (cat. no. 11733038, Thermo Fisher Scientific) and RNase/DNase-free PCR-grade water (Cat No. 10977015, Thermo Fisher Scientific), to a 5 μL total reaction volume. Thermocycling conditions comprised an initial 2 min denaturation at 95°C, followed by 45 cycles of 95°C for 15 sec and 60°C for 15 sec. All samples were run in duplicate.

### Genome assembly and annotation

Assemblies of the seven Vietnamese *N. seriolae* genomes were constructed from the quality-filtered sequence data using the Microbial Genome Assembly Pipeline (MGAP) v1.1 (https://github.com/dsarov/MGAP---Microbial-Genome-Assembler-Pipeline) and EM150506 (GenBank reference CP017839.1) as the scaffolding reference. MGAP wraps Trimmomatic (Bolger et al., 2014), Velvet (Zerbino and Birney, 2008), VelvetOptimiser (https://github.com/tseemann/VelvetOptimiser), ABACAS (Assefa et al., 2009), IMAGE (Tsai et al., 2010), SSPACE (Boetzer et al., 2011, Boetzer et al., 2010), GapFiller (Boetzer and Pirovano, 2012, Nadalin et al., 2012), and Pilon (Walker et al., 2014) into a single tool. Assemblies were primarily annotated using the Rapid Annotations using Subsystems Technology (RAST) server v2.0 with SEED data by default features (RAST annotation scheme: RASTtk, automatically fix errors, fix frameshifts, build metabolic model, backfill gaps, turn on debug, verbose level: 0, and disable replication: yes). RAST was also used to group genes into functional subsystems (akin to Clusters of Orthologous Groups). Annotated genomes were then compared with results provided by Prokka v1.8 (Seemann, 2014). In cases where aberrant results arose between the two tools, the functional prediction of RAST was checked and manually corrected by using BLASTP to search for similar proteins in the UniProtKB database (http://www.uniprot.org/blast/). The clustered regularly interspaced short palindromic repeat (CRISPR)/Cas region finder program (https://crisprcas.i2bc.paris-saclay.fr) was used to identify regular repeats and the intervening spacer sequences (Couvin et al., 2018). The assembled genomes for all Vietnamese strains are available from NCBI under BioProject PRJNA551736 (Table 3 3).

**Table 3.** Genetic and genomic features of the Vietnamese *Nocardia seriolae* strains compared with the South Korean EM150506 strain according to RAST

| Strains\Feature | Genome size (Mbp) | Level of completion | Sequencing platform | Sequencing depth | GC% | N50 (bp) | L50 (bp) | Total no. proteins | No. RNA | No. hypothetical proteins | No. proteins with function prediction | No. proteins assigned to subsystem | NCBI accession no. |
| --- | --- | --- | --- | --- | --- | --- | --- | --- | --- | --- | --- | --- | --- |
| KH_11 | 7.66 | Draft | NextSeq 500 | 340X | 68.3 | 27077 | 90 | 7655 | 58 | 3560 | 4465 | 2055 | WMKE000000000.1 |
| KH_21 | 7.72 | Draft | NextSeq 500 | 424X | 68.2 | 42752 | 58 | 7657 | 66 | 3597 | 4428 | 2033 | WMKF000000000.1 |
| NT_50 | 7.96 | Draft | NextSeq 500 | 395X | 68.2 | 29134 | 86 | 7640 | 66 | 3571 | 4437 | 2063 | WMKG000000000.1 |
| PY_31 | 7.68 | Draft | NextSeq 500 | 408X | 68.3 | 40217 | 62 | 7602 | 62 | 3212 | 4818 | 2220 | WMKC000000000.1 |
| PY_37 | 7.55 | Draft | NextSeq 500 | 326X | 68.3 | 19107 | 126 | 7707 | 51 | 3549 | 4525 | 2087 | WMKD000000000.1 |
| VT_45 | 7.94 | Draft | NextSeq 500 | 404X | 68.2 | 33835 | 70 | 7958 | 67 | 3609 | 4718 | 2054 | WMKB000000000.1 |
| VT_62 | 7.70 | Draft | NextSeq 500 | 433X | 68.3 | 40217 | 62 | 7643 | 63 | 3580 | 4428 | 2052 | WMKH000000000.1 |
| UTF1 | 8.12 | Complete | PacBio | 133X | 68.1 | 8121733 | 1 | 7890 | 75 | 3572 | 4683 | 2219 | AP017900.1 |
| U-1 | 7.77 | Draft | Roche 454; MiSeq | 179X | 68.3 | 42866 | 56 | 7757 | 69 | 3645 | 4497 | 2291 | BBYQ000000000.1 |
| N-2927 | 7.76 | Draft | Roche 454 | 160X | 68.3 | 45841 | 54 | 7627 | 66 | 3225 | 4841 | 2245 | BAWD000000000.2 |
| NBRC15557 | 7.61 | Draft | Roche 454; HiSeq 1000 | 112X | 68.3 | 45757 | 51 | 7527 | 64 | 3190 | 4768 | 2211 | NZ_BJWY01000001.1 |
| SY-24 | 7.89 | Draft | MiSeq | 100X | 68.2 | 46867 | 52 | 7632 | 66 | 3227 | 4845 | 2230 | MVAC000000000.1 |
| NK201610020 | 8.31 | Complete | HiSeq; PacBio | 100X | 68.1 | 4999276 | 1 | 8133 | 78 | 3398 | 5185 | 2306 | NZ_CP063662.1 |
| HSY-NS01 | 7.91 | Draft | HiSeq | 126X | 68.2 | 50962 | 50 | 7947 | 70 | 3727 | 4605 | 2133 | PXZE000000000.1 |
| HSY-NS02 | 7.76 | Draft | HiSeq | 110X | 68.2 | 46515 | 51 | 7801 | 69 | 3301 | 4932 | 2225 | RCNK000000000.1 |
| ZJ0503 | 7.71 | Draft | MiSeq | 100X | 68.3 | 46136 | 50 | 7579 | 66 | 3212 | 4798 | 2204 | JNCT000000000.1 |
| CK-14008 | 8.37 | Draft | PacBio | 139X | 68.1 | 8263617 | 1 | 8212 | 78 | 3422 | 5244 | 2347 | MOYO000000000.1 |
| MH196537 | 8.26 | Complete | PacBio | 118X | 68.1 | 8262437 | 1 | 8074 | 78 | 3368 | 5155 | 2296 | CP059737.1 |
| EM150506 | 8.30 | Complete | PacBio | 156X | 68.1 | 8304518 | 1 | 8068 | 77 | 3338 | 5175 | 2277 | CP017839.1 |

### Virulence and Antimicrobial Resistance Profile Determination

The identification of antimicrobial resistance-and virulence-related genes among the Vietnamese *N. seriolae* genomes were performed using RAST and the Virulence Factor Database (VFDB), Victors, and PATRIC Virulence Factor (VF) databases available on the Pathosystems Resource Integration Center (PATRIC) (Aziz et al., 2008, Wattam et al., 2013). In addition, homologues of experimentally verified pathogenicity determinants within other members of the *Nocardia* genus were searched for in the *N. seriolae* genomes.

## Results

### PFGE genotypes

Twenty *N. seriolae* isolates from four Vietnamese coastal provinces (Fig.1) were subjected to *Xba*I and *Ase*I digestion to determine isolate relatedness across provinces. Restriction fragment sizes ranged from 40kb-1.1Mbp. PFGE with *Xba*I alone resulted in between 19 and 21 restriction fragments among the Vietnamese strains; similarly, between 16 and 20 fragments were identified using *Ase*I. Seven distinct patterns (labelled as pulsotypes NsX1-NsX7) were present using *Xba*I­digested DNA fragments, and ten patterns (labelled as pulsotypes NsA1-NsA10) for *Ase*I. Using the ≥80% similarity cut-off and ‘fewer than six bands of difference’ Tenover criteria, only one cluster was identified for each enzyme (Tenover et al., 1995a, Calvez et al., 2015a). Even when combining data from both enzymes, the 20 Vietnamese isolates were still closely related, irrespective of their geographic origin, as shown by their categorisation into a single cluster that was distinct from the Japanese type strain (Fig. 3).

**Fig. 3.**
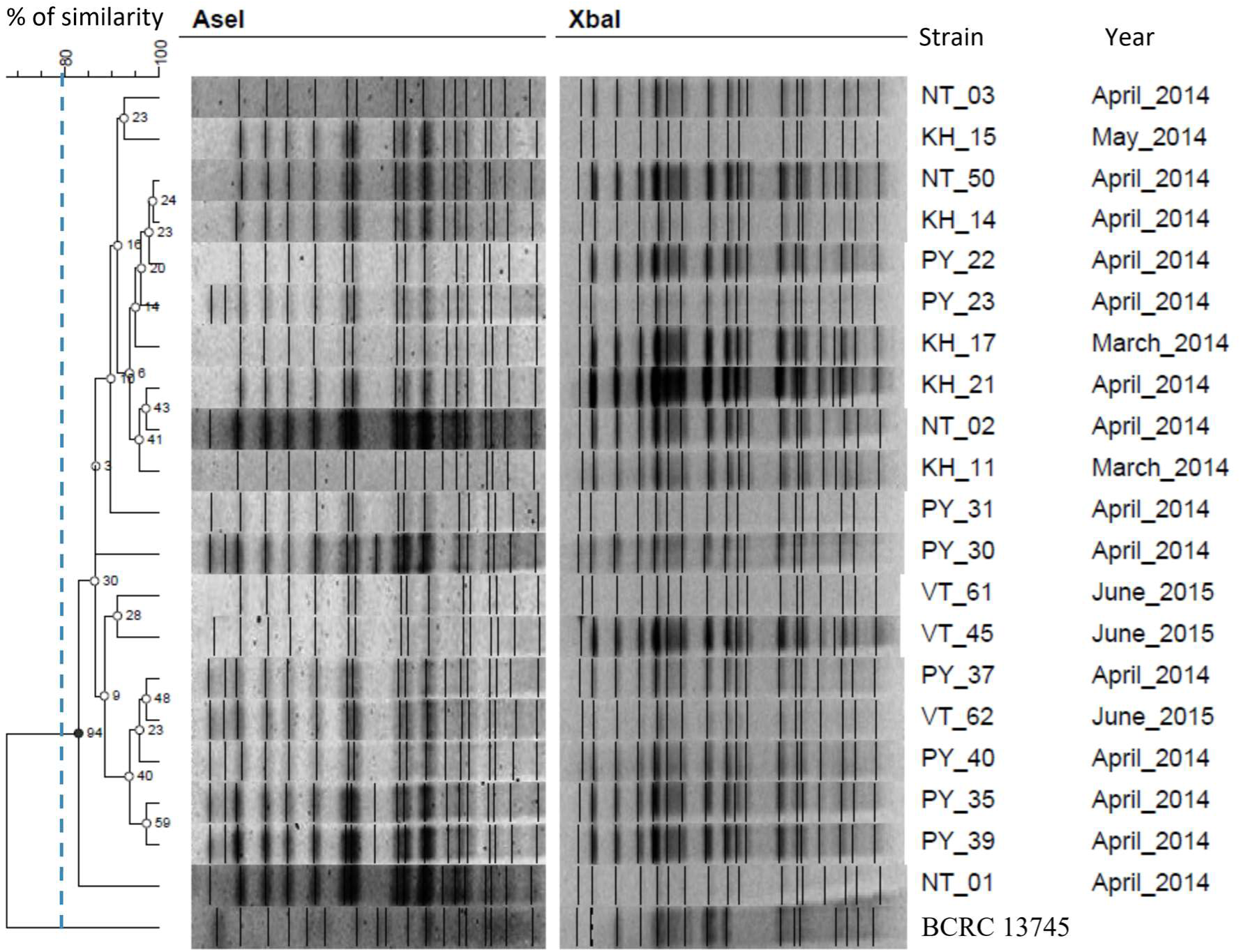
Pulsed-field gel electrophoresis dendrogram of *Ase*I and *Xba*I-digested genomic DNA from 20 representative *Nocardia seriolae* strains collected in four Vietnamese provinces. A type strain, BCRC 13745 (Japan), was included for comparison. Cluster analysis of genetic distances was performed using the Dice coefficient and UPGMA method (tolerance and optimisation 1%). Two pulsotypes were identified based on an 80% similarity cut-off. Numbers at tree nodes indicate the percentage of replicate trees in which the same clusters were found after 1,000 bootstrap replicates.

### Phylogenomic analysis

Based on the PFGE results, seven geographically diverse Vietnamese isolates were Illumina-sequenced, resulting in high-coverage draft genomes (Table 3). These genomic data were generated to address two questions: i) whether comparative genomics, like PFGE, would reveal minimal genetic diversity among the Vietnamese *N. seriolae* strains, and: ii) whether phylogenomic analysis could identify a potential origin for nocardiosis in Vietnamese aquaculture facilities. The seven Vietnamese genomes generated in this study, plus the sequences of 12 publicly available *N. seriolae* strains (all from other Asian countries), were compared to identify phylogenetically informative SNPs. A total of 8,206 SNPs were identified; 7,517 (91.6%) were located in coding regions and comprised 126 nonsense, 5,163 missense, and 1,531 silent variants. Of the 8,206 SNPs, 7,275 high-confidence, orthologous, core genome, biallelic SNPs were identified among the 19 *N. seriolae* strains; these SNPs were used for phylogenomic reconstruction.

The phylogenomic dendrogram revealed five distinct strain clusters (Fig. 4). As with PFGE, the seven Vietnamese isolates were highly clonal, with all strains clustering into a single unique ‘Vietnamese’ clade. Within this clade were two subclades that differed by 103 SNPs. These subclade SNPs were well-distributed across the genome, with no evidence of SNP clusters due to recombination. The phylogenomic analysis also suggested that *N. seriolae* undergoes very little, if any, recombination, as demonstrated by a very high consistency index of 0.997; in other words, homoplastic SNP characters, which are more common following recombination events (Crispell et al., 2019), were essentially absent. Within the two Vietnamese subclades, isolates were virtually identical (0-2 SNPs), indicating limited genomic alterations among these lineages (Fig. 4). Notably, there was no link between geographic region and subclade placement, with strains from Phú Yên, Khánh Hòa, and Vũng Tàu falling into both Vietnamese subclades, indicating frequent *N. seriolae* transmission events between regions. The most recent common ancestor of the Vietnamese strains differed by 217 SNPs from the next closest known strain, UTF1, which was isolated from cultured yellowtail that succumbed to nocardiosis in 2008 in Miyazaki Prefecture, Japan (Yasuike et al., 2017).

**Fig. 4.**
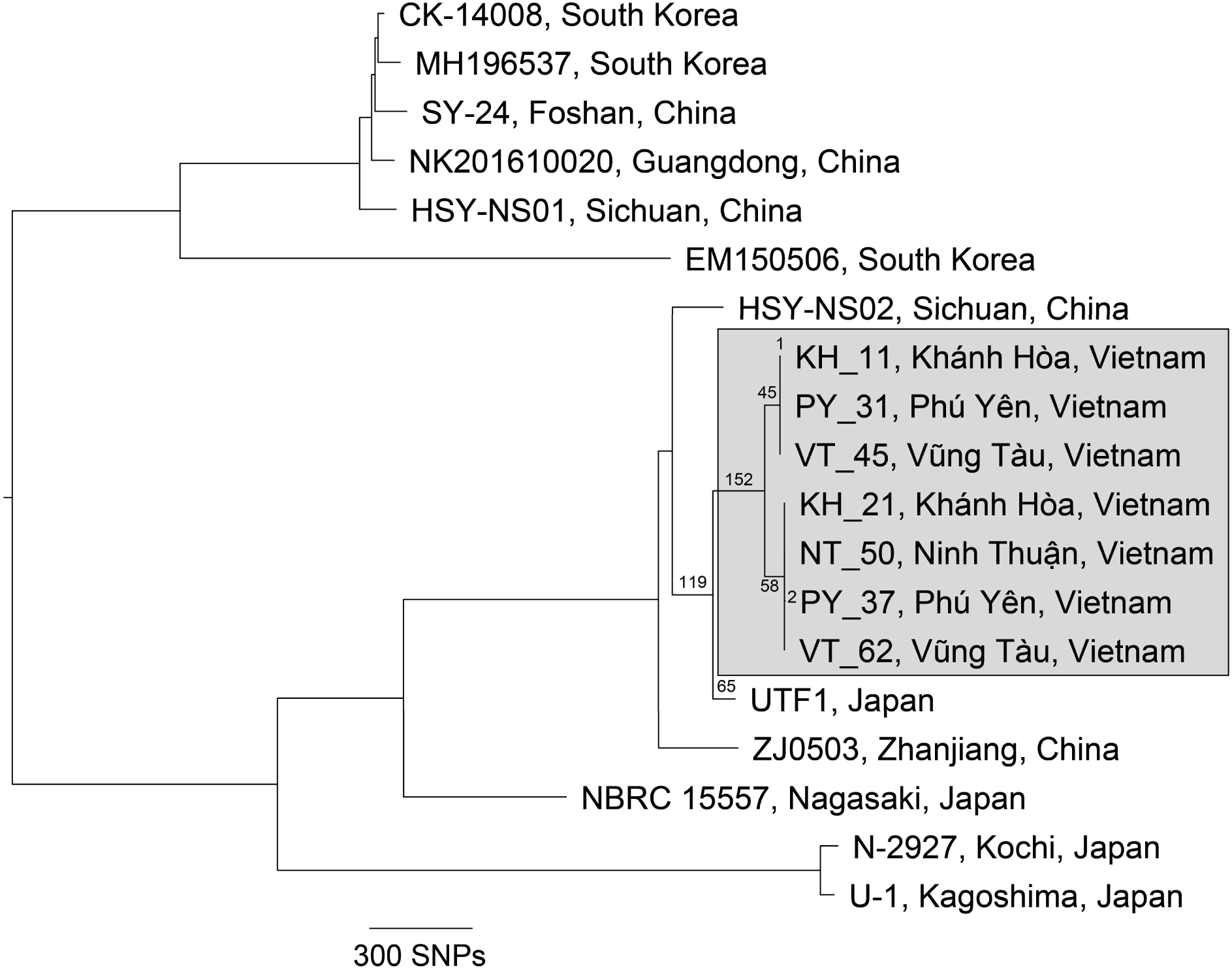
Midpoint-rooted maximum parsimony phylogenomic analysis of seven Vietnamese (KH_11, KH_21, NT_50, PY_31, PY_37, VT_62, and VT_45; grey box) and 12 non-Vietnamese *Nocardia seriolae* genomes. A total of 7,275 high-confidence biallelic, orthologous, core-genome single-nucleotide polymorphisms (SNPs) were used to construct the phylogeny. Branch lengths within the Vietnamese clade are labelled and refer to the number of SNPs along each branch. Consistency index=0.997.

### SNP genotyping

SYBR-MAMA assays demonstrated clear distinction of SNP genotypes. For the SNP1 assay, the two Taiwanese strains amplified the non-Viet allele earlier than the Viet allele (ΔCt range: 2.8 to 5.5); in contrast, all Vietnamese strains amplified the Viet allele earlier than the non-Viet allele (ΔCt range: 6.0 to 9.3). For the SNP2 assay, 10 Vietnamese strains belonging to Clade 1 amplified the Clade 1 allele earlier than the Clade 2 allele (ΔCt range: 9.9 to 13.4), whereas 10 Clade 2 strains amplified the Clade 2 allele earlier (ΔCt range: 4.5 to 8.1) (Table 1). No amplification was observed for the no-template controls.

### Genome Assembly and Functional Annotation

To gain deeper insights into the seven Vietnamese *N. seriolae* genomes, we conducted a comparative analysis of genome assembly metrics and gene function. The Vietnamese genomes possess 6,937 core genes and encode 1-6 ribosomal RNA genes and 49-63 transfer RNA genes. Total assembly length ranged from 7.55 to 7.96 Mbp, smaller than the closed genomes EM150506 (8.30Mbp), MH196537 (8.26Mbp), UTF1 (8.12Mbp), and draft genomes reported for CK-14008 (8.37Mbp), NK201610020 (8.31Mbp), but similar to other draft genomes of this species (range: 7.61 to 7.91Mbp). GC content (68.2 to 68.3%) was comparable to previously sequenced *N. seriolae* genomes (Table 3 and Fig. 5).

**Fig. 5.**
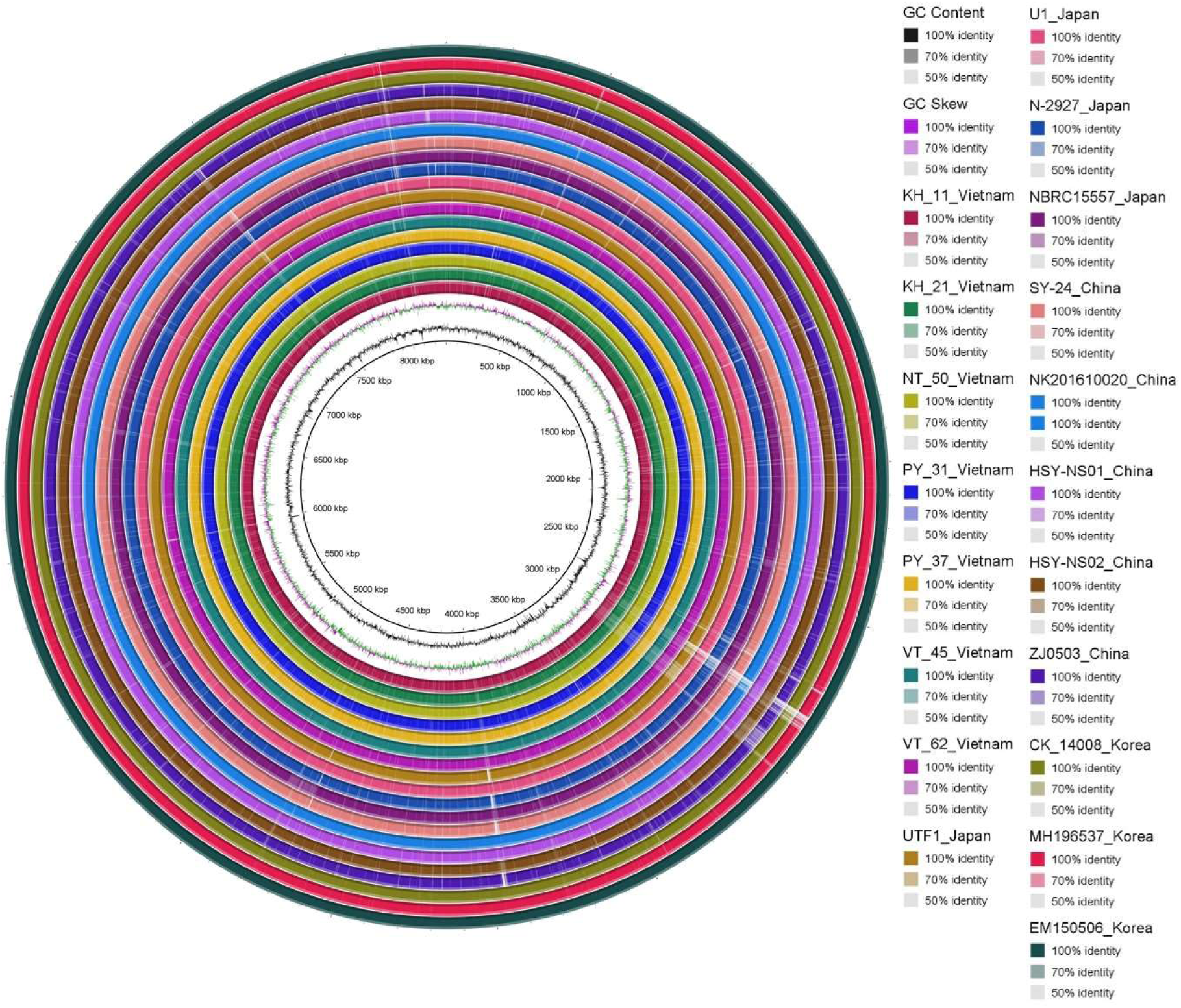
Whole-genome sequence comparison of *Nocardia seriolae* strains from Vietnam and other Asian countries against the EM150506 (South Korean) reference genome using the circular BLASTn alignment in BLAST Ring Image Generator (Alikhan et al., 2011). The innermost circle shows genome scale (bp), the black irregular ring represents %GC content, and the irregular purple/green ring represents GC skew. Outer colour rings (innermost first) represent Vietnamese strains (KH_11, KH_21, NT_50, PY_31, PY_37, VT_45, VT_62) and 12 strains from Japan, China, and South Korea. The outermost circle (dark green) represents the EM150506 reference genome.

RAST predicted between 7,602 and 7,958 coding DNA sequences in the Vietnamese *N. seriolae* genomes, of which 45.8% (range: 42.2-47.0%) are of unknown function (‘hypothetical proteins’). Of the 59.1% (range: 57.8-63.4%) coding DNA sequences with RAST function predictions, 45.8% (range: 43.5-50.9%) grouped into 308-330 functional subsystems belonging to 24 protein family categories. These predictions are similar to the previously reported *N. seriolae* genomes (Table 4). Little difference was found in the number of genes in family categories among Vietnamese vs. non-Vietnamese strains (Table 4). No plasmids were identified in any of the Vietnamese genomes, consistent with most *N. seriolae* genomes lacking plasmids; the only exception is CK-14008 from South Korea, which potentially harbours two plasmids (Han et al., 2018).

**Table 4.**
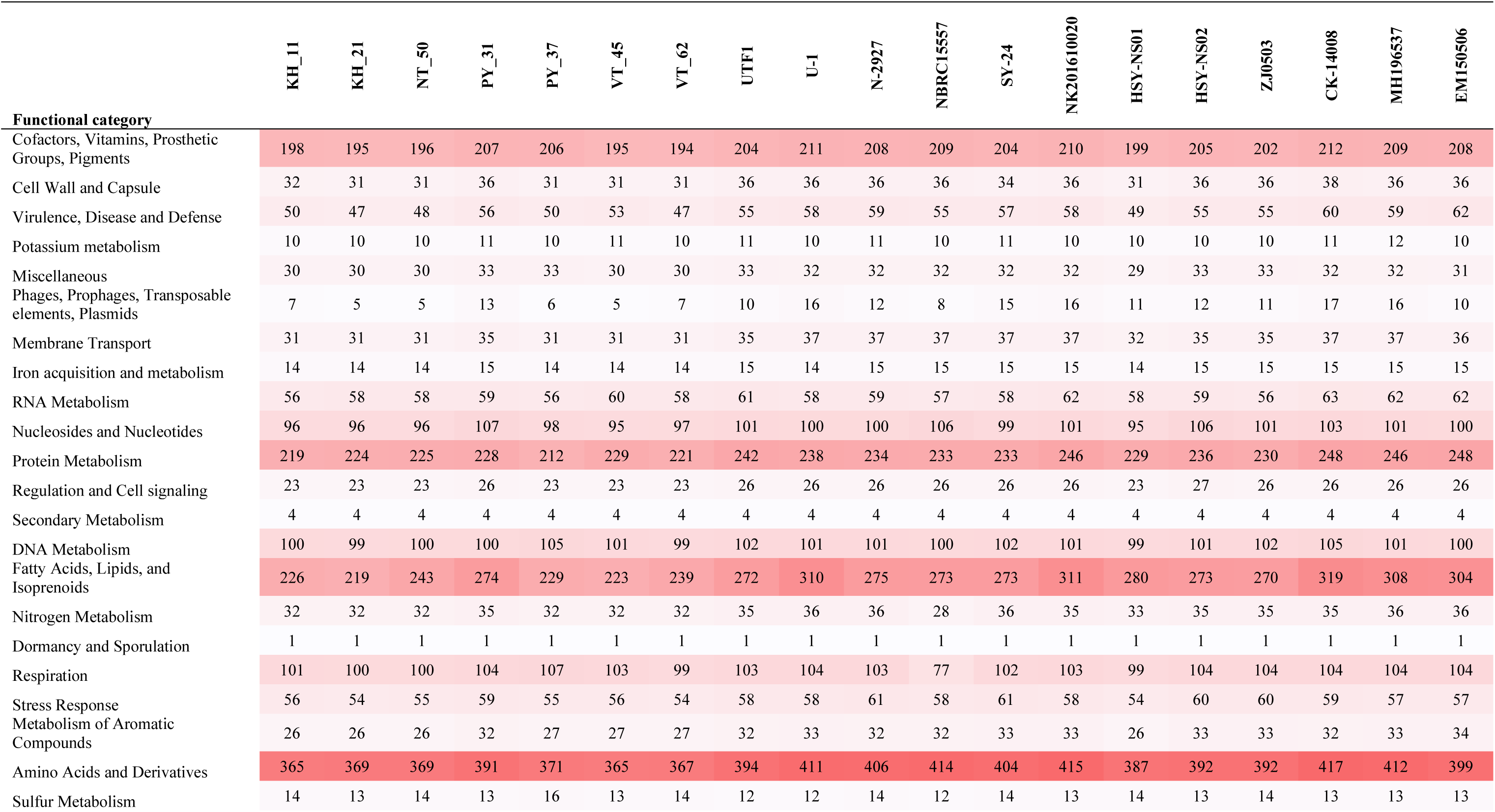
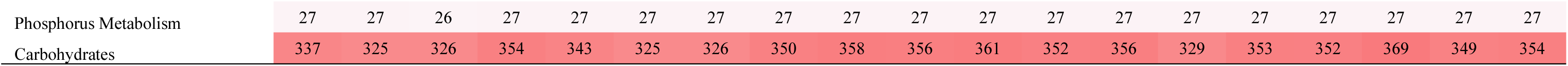
Number of genes for each *Nocardia seriolae* strain associated with the 24 general Clusters of Orthologous Groups functional categories predicted by RAST

Between three and six CRISPR arrays were found in the Vietnamese strains, with lengths varying from 73 to 114 bp. Each array is made of two direct repeats and one spacer without nearby Cas (CRISPR-associated) genes. Notably, the same CRISPR array structure was found in all 19 *N. seriolae* genomes (Data S2).

### Virulence and antimicrobial/toxin resistance profiles

To explore the pathogenic potential of the Vietnamese *N. seriolae* strains, we assessed their virulence and antimicrobial/toxin resistance gene content in comparison to non-Vietnamese genomes. RAST, VFDB, Victors, and VF databases found between 182 and 202 genes that encode virulence and resistance factors, including gene products associated with Adherence (*n*=50-54), Cellular metabolism & nutrient uptake (*n*=10), Damage (*n*=6-7), Invasion and intracellular survival (*n*=33-36), Resistance to antibiotics and toxic compounds (*n*=65-81), and Other (*n*=16-18) (Data S3). In general, virulence factors and antimicrobial/toxin resistance factors were almost identical in number among the Vietnamese strains and were comparable to non-Vietnamese strains. However, some genes were absent in most Vietnamese strains but present in most non-Vietnamese strains, such as “MCE-family protein Mce1D”, “MCE-family protein Mce1F”, “Chromate transport protein ChrA”, “NAD(P)H oxidoreductase YRKL (EC 1.6.99.-) Putative NADPH-quinone reductase (modulator of drug activity B) Flavodoxin 2”, and “Tellurite resistance protein TerB”. In contrast, “Hemolysins and related proteins containing cystathionine-β-synthase domains” was found only in EM150506. Several experimentally verified virulence factors identified in *N. seriolae* and other *Nocardia* spp., including catalase, superoxide dismutase, phospholipase C, and protease (Vera-Cabrera et al., 2013), were present in all Vietnamese and non-Vietnamese strains, indicating that they are highly conserved genes within this genus.

## Discussion

*N. seriolae* is an emerging global aquaculture pathogen that has caused devastating fish outbreaks and mass fish mortalities in recent decades, particularly in Asia and the Americas. The presence of this bacterium in fish farms requires close surveillance; its control at present solely relies on antimicrobial agents. Analysis of *N. seriolae* genetic diversity, pathogenic and resistance potential, and population structure is essential for understanding the origin, dissemination, and antimicrobial susceptibility potential of this economically important pathogen, which will, in turn, inform better farm management practices and limit accidental transmission into naïve fish populations.

*N. seriolae* caused severe mortalities in fish farms in several Vietnamese provinces between 2010 and 2015, less than a decade after the first *T. falcatus* fingerlings were imported from China and Taiwan. To better understand the genetic diversity and putative origin of the Vietnamese outbreak, we employed PFGE and WGS to examine strains obtained from diseased fish from four Vietnamese coastal provinces in 2014 and 2015. PFGE has conventionally been considered the “gold standard” for studying the genetic diversity of many different pathogenic bacteria species, including *N. seriolae* (Shimahara et al., 2008, Shimahara et al., 2009, Calvez et al., 2015b, Sun et al., 2016). PFGE has previously identified multiple pulsotypes among isolates retrieved from fish in Japan and Taiwan (Shimahara et al., 2008, Shimahara et al., 2009). Notably, one study identified identical pulsotypes between certain Taiwanese 1997-2007 outbreak strains and Japanese *N. seriolae* isolated from yellowtail in 2002 (pulsotypes X1 and A1) and 2005 (pulsotype X11) (Shimahara et al. (2009), suggesting at least two transmission events between Taiwan and Japan. Unlike *N. seriolae* from Japan and Taiwan, all 20 Vietnamese isolates fell into a single cluster, even when using a combination of *Xba*I and *Ase*I. However, PFGE lacked the resolution to differentiate Vietnamese isolates into the two clades identified using phylogenomic analysis. This limited resolution has also been documented for other bacteria such as *Salmonella enterica* (den Bakker et al., 2011)*, Listeria monocytogenes* (Kwong et al., 2016), and *Escherichia coli* (Lee et al., 2017). It was unfortunately not practical to compare the Vietnamese pulsotypes with published studies due to known challenges with interlaboratory standardisation using PFGE (Seifert et al., 2005); therefore, it is not known whether the Vietnamese PFGE cluster has been previously reported.

Next-generation sequencing provides excellent resolution, accuracy, and data portability, and as such, has begun replacing PFGE as the new gold standard for nocardiosis outbreak analyses (Uelze et al., 2020). To illustrate the value of WGS for nocardiosis epidemiological investigations, we sequenced seven representative Vietnamese *N. seriolae* strains and compared them with all publicly available genomes available at the time (*n*=12). Like PFGE, the limited genomic variation (0-2 SNPs) observed among Vietnamese strains confirms a recent, single introduction into Vietnam, with subsequent dissemination across multiple mariculture facilities within the South-Central Coast region. Phylogenomic analysis showed that Vietnamese strains were most closely related to UTF1, isolated from farmed yellowtail in Japan in 2008 (Yasuike et al., 2017); this strain differed from the Vietnamese common ancestor by just 217 SNPs. Shimahara et al. (2009) have previously postulated that transboundary translocation of live fish stocks asymptomatically infected with *N. seriolae* from China and Hong Kong may have introduced new strains into Japan. Based on our genomic analysis, it is also plausible that *N. seriolae* from Japan has been introduced into other countries such as Vietnam given that international export of valuable aquaculture fish species is not uncommon; however, there is a paucity of information about import-export of live fish stocks from Japan or Vietnam, and as such, this hypothesis cannot be confirmed.

Whilst our results suggest a likely Asian origin for the Vietnamese outbreaks, there are few publicly available *N. seriolae* genomes (only 20 as of 01Nov21, including seven from our study), and none from other Asian regions such as Taiwan (Shimahara et al., 2009), Singapore, Malaysia, Indonesia (Labrie et al., 2008) or non-Asian regions such as Mexico (Del Rio-Rodriguez RE, 2021) and USA (Cornwell et al., 2011) where *N. seriolae* outbreaks have been documented; therefore, the precise origin of the Vietnamese outbreaks and mode of *N. seriolae* introduction currently remains unresolved. Concerningly, our results, and those of others, demonstrate that, unchecked, *N. seriolae* transmission may represent a substantial unmitigated risk to fish aquaculture. It is thus an utmost imperative to establish domestic and international monitoring processes for *N. seriolae* for both farmed and wild species, including the implementation of molecular methods to characterise new outbreaks, to prevent the spread of this devastating pathogen into new environments, and associated heavy economic losses and food security concerns.

To facilitate the rapid identification of *N. seriolae* genotypes among our Vietnamese strains, we designed inexpensive SYBR-MAMA assays targeting two phylogenetically informative SNPs. The first SNP assay robustly differentiates Viet from non-Vietnamese strains, thereby permitting prospective identification of newly transmitted strains into Vietnam, an essential facet in future fish importation biocontrol efforts. This assay can also be used to monitor for the emergence of Vietnamese strains in new regions, such as new aquaculture facilities in Vietnam, or prior to export of fingerlings to other countries. The second SNP assay rapidly differentiates strains belonging to the two Vietnamese clades. By applying this second assay to the 20 Vietnamese strains, we observed that both clades were well-disseminated across all four provinces: Khánh Hòa, Ninh Thuận, Phú Yên, and Vũng Tàu. Phylogenomic analysis of seven representative Vietnamese strains also showed dispersal of these two clades among three of the four provinces. Although unconfirmed, it is probable that the widespread trade of eggs, fingerlings, and live permit for aquaculture in Vietnam since industry inception in the early 2000s, including local unmonitored trade among fish farmers, has driven the successful dissemination of *N. seriolae* among Vietnamese permit farms. Taken together, our findings highlight the large risk of undetected *N. seriolae* dispersal among mariculture facilities and the need for establishing strict monitoring practices to prevent further pathogen transmission.

WGS is currently laborious, expensive, and inaccessible to most laboratories in Vietnam and many other Asian countries. Using comparative genomics, we established a catalogue of SNPs specific to each clade and subclade. This SNP database may be useful for both targeted resequencing efforts and the design of phylogenetically robust genotyping methods to permit source tracing of future *N. seriolae* outbreaks without the requirement for further WGS or bioinformatic analyses. The SYBR-MAMA assays developed in this study successfully detected two phylogenetically informative SNPs, with genotyping results fully concordant with WGS, confirming that SYBR-MAMA is a valuable and inexpensive diagnostic method for SNP characterisation.

Very little is known about the pathogenesis of *Nocardia* spp., which are capable of invading host macrophages and preventing the fusion of phagosomes with lysosomes, leading to long-term survival and proliferation in host cells (Davis-Scibienski and Beaman, 1980). Due to the paucity of available genomic data for this pathogen, a final aspect of this study was to better understand virulence and antimicrobial resistance factors encoded by the *N. seriolae* genome. Our analysis of 19 *N. seriolae* genomes is the largest genomic assessment of this pathogen to date, and largely corroborates the conclusions drawn from a previous analysis of seven *N. seriolae* genomes, which showed that *N. seriolae* have >99.9% Orthologous Average Nucleotide Identity values (Han et al., 2018). More than 180 genes were found to encode for antimicrobial resistance and virulence factors in the Vietnamese strains. We catalogued the 180 core (present in all strains) genes, including genes associated with Adherence (*n*=49), Cellular metabolism & nutrient uptake (*n*=10), Damage (*n*=6), Invasion and intracellular survival (*n*=33), Resistance to antibiotics and toxic compounds (*n*=26), and Others (*n*=11) that may possibly account for the main virulence traits of this fish pathogen. Analysis of the genome content of seven Vietnamese *N. seriolae* strains revealed that, like non-Viet strains, they encode a high proportion of ‘hypothetical protein’ genes (i.e. 45.8%), a finding that highlights the need for more studies to investigate the functions of these genes. The presence of conserved genes encoding β-lactamase class C-like and penicillin-binding proteins (*n*=11), multidrug resistance protein ErmB (*n*=1), probable multidrug resistance protein NorM (*n*=1), and a small multidrug resistance family protein (*n*=1) in all *N. seriolae* genomes, may explain observed antimicrobial resistance towards penicillin and cephalexin, two β-lactam antibiotics that are commonly used to treat nocardiosis in Vietnamese permit fish farms (data not shown).

In conclusion, our study provides novel insights into the epidemiology of *N. seriolae* outbreaks in farmed permit fish farm in Vietnam. Our detailed molecular and genomic analyses revealed minimal genomic diversity among Vietnamese *N. seriolae* isolates; unlike PFGE, WGS detected strain variation at single-base resolution, and identified two distinct Vietnamese clades that share recent ancestry. Our results indicate recent importation of a single *N. seriolae* clone into Vietnam, which led to a nationwide outbreak of nocardiosis in permit fish farms. The analysis of additional genomes, particularly from more geographic regions, will be important for better understanding *N. seriolae* evolution, and will enable more precise investigations into the origin and transmission of this devastating pathogen. Finally, our SNP assays provide a rapid and inexpensive method for genotyping of ongoing and future nocardiosis outbreaks in Vietnam.

## Author contributions

CL: Project design, sample collection, sample and data analysis, results interpretation, drafting paper.

DSS: Data analyses and interpretation, drafting and revising paper.

EPP: Supervision, data analyses and interpretation, drafting and revising paper.

TTAN: Assistance in the sample preparation and drafting paper.

DP: Sample collection guidance, drafting and revising paper.

HV-K: Sample collection guidance, drafting and revising paper.

IK: Assisting with the project design, revising paper.

WK: Supervision, advising on project design, drafting paper.

S-CC: Assistance in PFGE analyses, drafting paper.

MK: Supervision, project design, revising paper.

All authors read and approved the final manuscript.

## Acknowledgments

We gratefully acknowledge the financial support and laboratory facilities provided by the Genecology Research Centre, the University of the Sunshine Coast, and Nha Trang University. This research was supported by an Australia Awards PhD scholarship to CL, which is funded by the Australian Department of Foreign Affairs and Trade. DSS and EPP were supported by Advance Queensland fellowships (AQRF13016-17RD2 and AQIRF0362018, respectively).

